# Taxonomic context and genomic architecture jointly shape expression divergence across animals

**DOI:** 10.64898/2025.12.18.695241

**Authors:** Perry A. LaBoone, Antara Anika Piya, Raquel Assis

## Abstract

Gene expression divergence is a major source of phenotypic variation, yet the factors that shift regulatory optima remain incompletely understood. In particular, it is unclear how broad taxonomic differences and local genomic architecture interact to shape the tempo and mode of expression evolution. Here, we analyze single-copy orthologs between species pairs in *Drosophila* and mammals to quantify taxon-specific rates and magnitudes of expression divergence, assess the influence of genome architecture, and evaluate tissue-specific and functional patterns. Using a computational framework that predicts expression divergence and species-specific expression optima, we identify markedly higher divergence rates in *Drosophila* than in mammals, consistent with theoretical expectations and prior empirical work. Contrary to expectations, genes located in nested structures, in which one gene lies entirely within the intron of another, were no more likely to diverge than unnested genes in either taxon. However, when divergence occurred in *Drosophila*, nested genes exhibited larger shifts in expression optima, with the strongest effects among internal genes. Divergence rates were also higher among young than old nested genes in both taxa, although magnitudes of expression shifts were indistinguishable, suggesting that nesting contributes to early regulatory instability but does not typically trigger large regulatory changes. Tissue-level and functional analyses further revealed taxon-and architecture-specific signatures of expression divergence, including contrasting patterns across reproductive and neural tissues, as well as enrichment of core regulatory processes among unnested genes and enrichment of drug and xenobiotic metabolism among nested genes in *Drosophila*. Collectively, these findings demonstrate that taxonomic context and genome architecture shape expression evolution in distinct and measurable ways, giving rise to contrasting patterns of regulatory divergence.

## Introduction

Understanding how gene expression evolves between species is central to explaining phenotypic diversity and the mechanisms that generate new traits. Expression levels of orthologous genes often diverge substantially over evolutionary time, reflecting a mixture of neutral drift, compensatory changes, and lineage-specific selection (Khaitovich et al., 2004, 2006; Brawand et al., 2011; Fraser, 2011; Perry et al., 2012; Wittkopp and Kalay, 2012; Coolon et al., 2014; Signor and Nuzhdin, 2018; Cardoso-Moreira et al., 2019). Comparative transcriptomic studies across taxonomic groups consistently show that expression divergence varies widely among genes and tissues, and that this heterogeneity carries functional and evolutionary significance (Khaitovich et al., 2006; Brawand et al., 2011; Perry et al., 2012; Coolon et al., 2014; Cardoso-Moreira et al., 2019). Yet the factors that determine which genes maintain conserved expression profiles and which diverge rapidly remain unclear.

One axis along which divergence patterns differ is taxonomic context. For example, *Drosophila* and mammals often vary markedly in genome size and structure, effective population size, and regulatory architecture, all of which can influence the tempo and mode of expression evolution (Beckenbach et al., 1993; Adams et al., 2000; Hahn and Wray, 2002; Lynch and Conery, 2003; Jensen and Bachtrog, 2011; modENCODE Consortium et al., 2010; Signor and Nuzhdin, 2018; Moore et al., 2020). Prior comparative studies suggest that genome-wide expression divergence may proceed differently in these clades (Brawand et al., 2011; Perry et al., 2012; Coolon et al., 2014; Cardoso-Moreira et al., 2019), but systematic cross-taxon analyses remain limited, particularly those incorporating tissue-specific patterns (Wray, 2007; Romero et al., 2012). Such contrasts are essential for interpreting lineage-specific signatures of divergence and for contextualizing more specialized genomic features.

A second dimension of heterogeneity arises from genome architecture. Structural features such as gene order, regulatory neighborhoods, chromatin states, and physical arrangements along chromosomes can influence the accessibility and coordination of regulatory elements (Hurst et al., 2002; modENCODE Consortium et al., 2010; Bell and Spector, 2011; Dixon et al., 2012; Kundaje et al., 2015; Rowley and Corces, 2018; Moore et al., 2020). One of the most striking architectural configurations is the nested gene structure, in which an internal gene resides entirely within the intron of another, external host gene (Henikoff et al., 1986; Veeramachaneni et al., 2004; Yu et al., 2005). Nested structures represent the most common form of protein-coding overlap and are widespread across metazoan genomes, comprising approximately 10% of genes in *Drosophila* and 5% of genes in mammals (Assis, 2016, 2021). Prior work indicates that nested genes often originate through relocation events that place genes into introns (Assis et al., 2008). Because relocations expose genes to new regulatory environments and, in the case of nested constructs, may introduce transcriptional interference (Da Lage et al., 2003; Yu et al., 2005; Crampton et al., 2006; Osato et al., 2007; Assis et al., 2008), they have been hypothesized to influence gene expression. However, a recent study examining a small subset of 102 relocated genes in *Drosophila* found that only a modest fraction ( 23%) showed evidence of expression divergence following relocation (Piya et al., 2023). This discrepancy raises a broader question: Do relocations generally spur expression divergence, and how do such changes compare with genome-wide background levels?

Adding to this puzzle, previous studies show that expression evolution of nested genes follows strikingly different trajectories in *Drosophila* and mammals (Assis, 2016, 2021). In *Drosophila*, nested genes tend to exhibit greater expression divergence than unnested genes, whereas in mammals the two groups are largely comparable. Yet despite these clade-specific differences, both groups exhibit a consistent pattern in which internal nested genes diverge more than their external hosts (Assis, 2016, 2021). This pattern suggests that transcriptional interference or regulatory conflict may impose selective pressures favoring either rapid divergence or swift loss of internal genes whose expression overlaps excessively with that of their external hosts. Though these findings highlight nested structures as a potentially important driver of regulatory evolution, their consequences for genome-wide patterns of expression divergence have not been systematically evaluated across taxa.

Here, we investigate how both taxonomic context and genome architecture, specifically nested structures, contribute to genome-wide patterns of expression divergence. To address these questions, we analyze single-copy orthologs between pairs of species in two deeply separated clades: *Drosophila* (*D. melanogaster* and *D. pseudoobscura* and mammals (human and mouse). We use the neural network PiXi to classify ortholog pairs as “conserved” or “diverged” based on their expression profiles and to estimate their expression optima in each species (Piya et al., 2023). From these estimates, we compare rates of expression divergence, differences in expression optima, tissue-specific patterns of divergence, and functional enrichment between unnested and nested genes within each taxon. These analyses provide a framework for understanding how taxonomic context and genome architecture together shape the evolution of gene expression.

## Results

### Prevalence of genome-wide expression divergence

To quantify genome-wide patterns of expression divergence, we analyzed normalized expression data collected from matched tissues within each species pair: female head, male head, carcass, ovary, accessory gland, and testis in *Drosophila*; and brain, lung, heart, liver, spleen, kidney, colon, and testis in mammals (see *Methods*). We focused on single-copy genes to ensure unambiguous orthology and avoid confounding from duplication, identifying 8,268 1:1 orthologs between *D. melanogaster* and *D. pseudoobscura* and 9,245 1:1 orthologs between human and mouse (see *Methods*). To evaluate the influence of genomic architecture on expression divergence, we partitioned these orthologs into two subsets: genes unnested in both species of a taxon, and genes nested in at least one of the species. In total, we considered 7,157 unnested and 1,111 nested orthologs in *Drosophila* and 8,725 unnested and 520 nested orthologs in mammals (see *Methods*). These orthologs and their associated expression data were used as input to the PiXi neural network framework (Piya et al., 2023), which classified each pair as “conserved” or “diverged” and estimated species-specific expression optima (see *Methods*).

We first evaluated taxon-specific levels of expression divergence by comparing the proportions of genes classified as “diverged” in each taxon. Across all genes, expression divergence was substantially more frequent in *Drosophila* (14.79%) than in mammals (3.97%; *P* = 1.97 × 10*^−^*^253^; see *Methods*). A similar contrast was observed for unnested genes, with 15.16% “diverged” in *Drosophila* and 3.97% in mammals (*P* = 1.88 × 10*^−^*^251^; see *Methods*). This pattern aligns with theoretical expectations arising from the larger effective population size of *Drosophila* relative to mammals (Beckenbach et al., 1993; Lynch and Conery, 2003; Jensen and Bachtrog, 2011), as well as with empirical evidence for elevated rates of adaptive protein-coding and regulatory sequence evolution (Britten, 1986; Moriyama, 1987; Carroll, 2005) and rapid, pervasive expression evolution (Coolon et al., 2014; Nourmohammad et al., 2017) in *Drosophila*.

Among nested genes, 12.42% were classified as “diverged” in *Drosophila*, a significantly smaller proportion than the 15.16% observed for unnested genes (*P* = 1.07 × 10*^−^*^2^; see *Methods*). In contrast, 4.04% of nested mammalian genes were classified as “diverged” (*P* = 6.39 × 10*^−^*^11^; see *Methods*), which was not significantly different from the 3.97% observed for unnested genes (*P* = 0.91; see *Methods*). Thus, nested genomic architecture was associated with a reduced likelihood of expression divergence in *Drosophila* and showed no detectable effect in mammals.

### Effects of genomic architecture on expression divergence

Our finding that gene nesting does not influence the rate of expression divergence in mammals is consistent with previous work showing that mammalian genes typically retain their expression profiles after nesting (Assis, 2021). However, in *Drosophila*, the reduced rate of expression divergence among nested genes initially appeared to conflict with an earlier report of elevated expression divergence following nesting in this taxon (Assis, 2016). To investigate this apparent discrepancy, we compared magnitudes of expression divergence, which we quantified as absolute differences between predicted expression optima, for unnested and nested genes classified as “diverged” in each taxon. Consistent with prior reports (Assis, 2016, 2021), nested genes exhibited larger shifts in expression optima than unnested genes in *Drosophila*, whereas no difference was observed in mammals (Figure 1, left).

**Figure 1:**
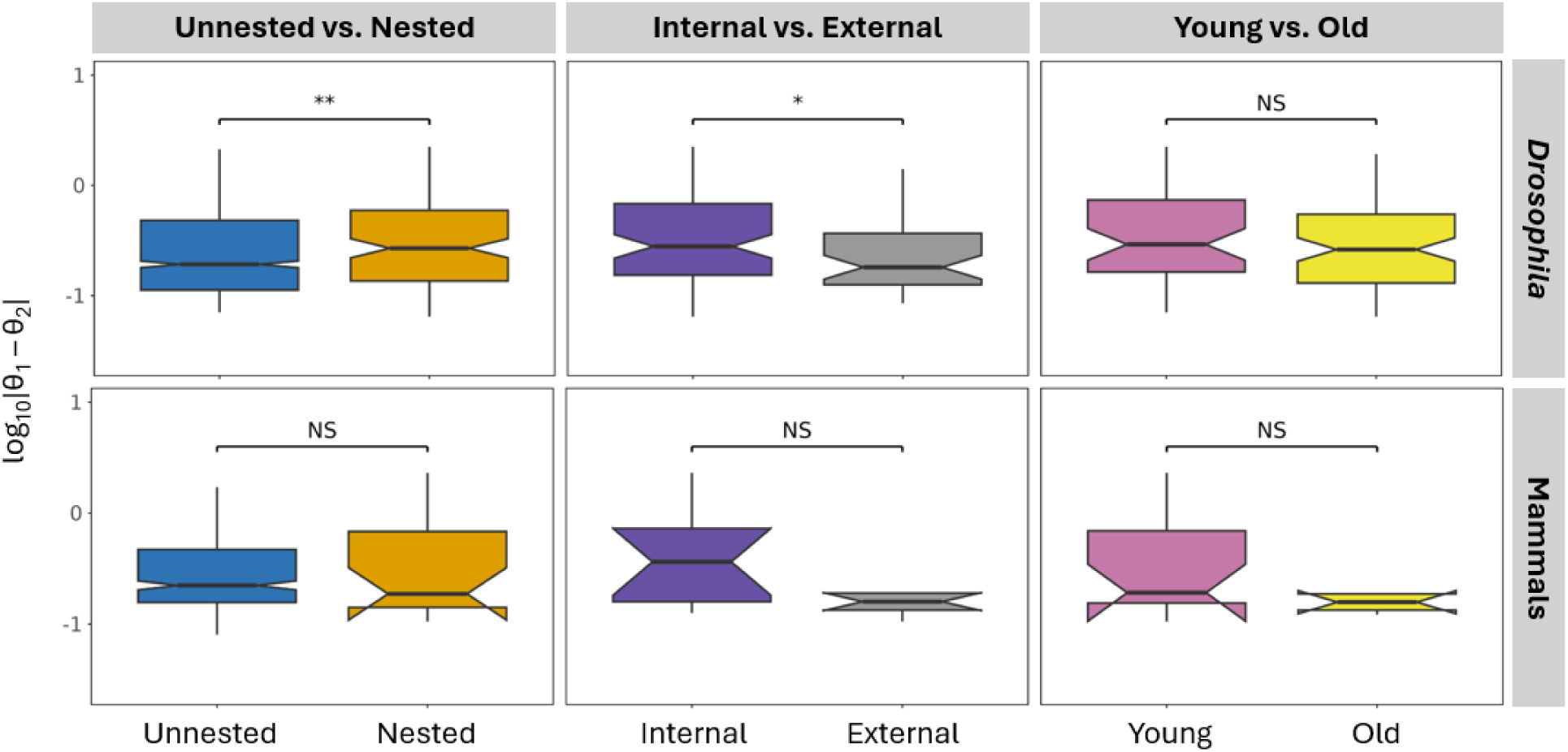
Magnitudes of expression divergence for unnested and nested orthologs. Box plots show distributions of absolute differences between species-specific expression optima for unnested and nested genes (left), internal and external nested genes (middle), and young and old nested genes (right) in *Drosophila* (top) and mammals (bottom). \**p <* 0.05, \*\**p <* 0.01, \*\*\**p <* 0.001, NS = not significant (see *Methods*).

We then asked whether internal and external nested genes contributed differently to these patterns. Our analysis revealed that expression divergence was significantly more frequent among internal (14.15%) than external (9.98%) nested genes in *Drosophila* (*P* = 8.07×10*^−^*^4^; see *Methods*), whereas in mammals, internal and external nested genes exhibited similar frequencies of divergence (4.44% vs. 3.60%; *P* = 0.41; see *Methods*). Internal genes also tended to display greater shifts in expression optima in *Drosophila*, but not in mammals (Figure 1, middle). These results indicate that although nested genes may not diverge more frequently than unnested genes in either taxon, the divergence that does occur in *Drosophila* is often pronounced and biased toward internal genes, consistent with prior work (Assis, 2016).

To examine whether the evolutionary age of nested structures influenced these outcomes, we compared rates of expression and shifts in expression optima between young and old nested genes. For this analysis, we defined a nested gene structure as young if it was present in only one species and old if it was shared by both (see *Methods*). In *Drosophila*, divergence rates were higher in young (14.48%) than in old (11.41%) nested genes, but this difference was not statistically significant (*P* = 0.07; see *Methods*). However, in mammals, divergence was significantly more frequent in young (5.21%) than in old (2.35%) nested genes (*P* = 3.49 × 10*^−^*^3^; see *Methods*). Despite these differences in rates, young and old nested genes did not differ in the magnitudes of their shifts in expression optima in either taxon (Figure 1, right). Together, these findings suggest that expression divergence is not generally driven by nesting itself, but rather reflects the selective retention of nesting events involving genes that were already sufficiently distinct.

### Functional characterization of expression divergence

To place expression divergence in a functional context, we first examined tissue-specific magnitudes of expression divergence for orthologous gene pairs classified as “diverged” in each taxon (Figure 2). Among *Drosophila* unnested genes, expression divergence was highest in carcass and consistently lowest across the three sex tissues (ovary, testis, and accessory gland). In mammals, divergence of unnested genes was highest in kidney and lowest in heart, lung, and colon. Tissue-specific patterns also differed between unnested and nested genes. In *Drosophila*, nested genes again showed the highest divergence in carcass but were followed by markedly elevated divergence in testis, approximately twofold higher than that observed for unnested genes; female and male head and accessory gland also exhibited nearly 1.5-fold higher divergence in nested relative to unnested genes. In mammals, nested genes displayed the highest divergence in brain and testis and the lowest divergence in lung and colon. Together, these results demonstrate that tissue-specific expression divergence varies across both taxa and genomic contexts.

**Figure 2:**
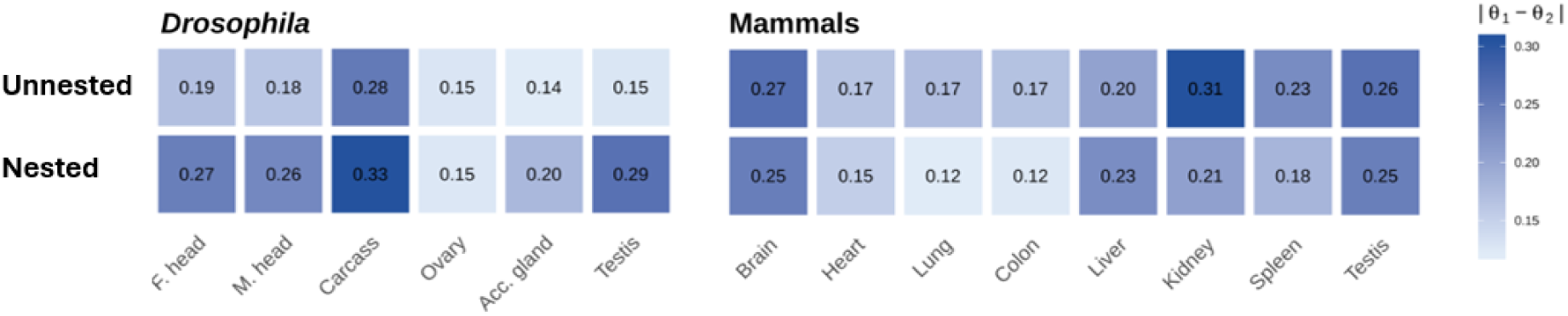
Magnitudes of tissue-specific expression divergence for unnested and nested orthologs in *Drosophila* and mammals. Heatmaps depict absolute differences between species-specific expression optima across individual tissues for unnested (top) and nested (bottom) genes in *Drosophila* (left) and mammals (right).

The observed tissue-specific differences prompted us to examine whether expression divergence is associated with particular biological functions. To address this question, we performed functional enrichment analyses of “diverged” genes in each taxon. To capture a broad range of functional categories, we incorporated annotations from the Gene Ontology (GO) Consortium (The Gene Ontology Consortium, 2021), curated pathways from the Kyoto Encyclopedia of Genes and Genomes (KEGG) (Kanehisa et al., 2016), signaling and regulatory networks from BioCarta (BioCarta, 2025), and protein domain features from InterPro (Blum et al., 2025) and the Simple Modular Architecture Research Tool (SMART) (Letunic and Bork, 2018). Enrichment was assessed by testing whether each target set of “diverged” genes was overrepresented in specific functional categories relative to the genome-wide background (see *Methods*).

Our functional enrichment analyses revealed distinct taxon- and architecture-dependent patterns. Among *Drosophila* “diverged” unnested genes, we observed significant enrichment for transcriptional regulation and RNA processing (Table S1), whereas mammalian “diverged” unnested genes were enriched primarily for cellular component terms related to mitochondrial and membrane localization (Table S2). In contrast, “diverged” nested genes in *Drosophila* exhibited enrichment for drug and xenobiotic metabolism, particularly cytochrome P450-mediated processes (Coelho et al., 2015) (Table S3), while no statistically significant functional enrichment was detected among mammalian “diverged” nested genes. Visualization of the top 15 enriched terms clarifies the architectural contrast within *Drosophila*, illustrating that nested genes exhibit fewer but substantially larger enrichment signals than unnested genes (Figure 3).

**Figure 3:**
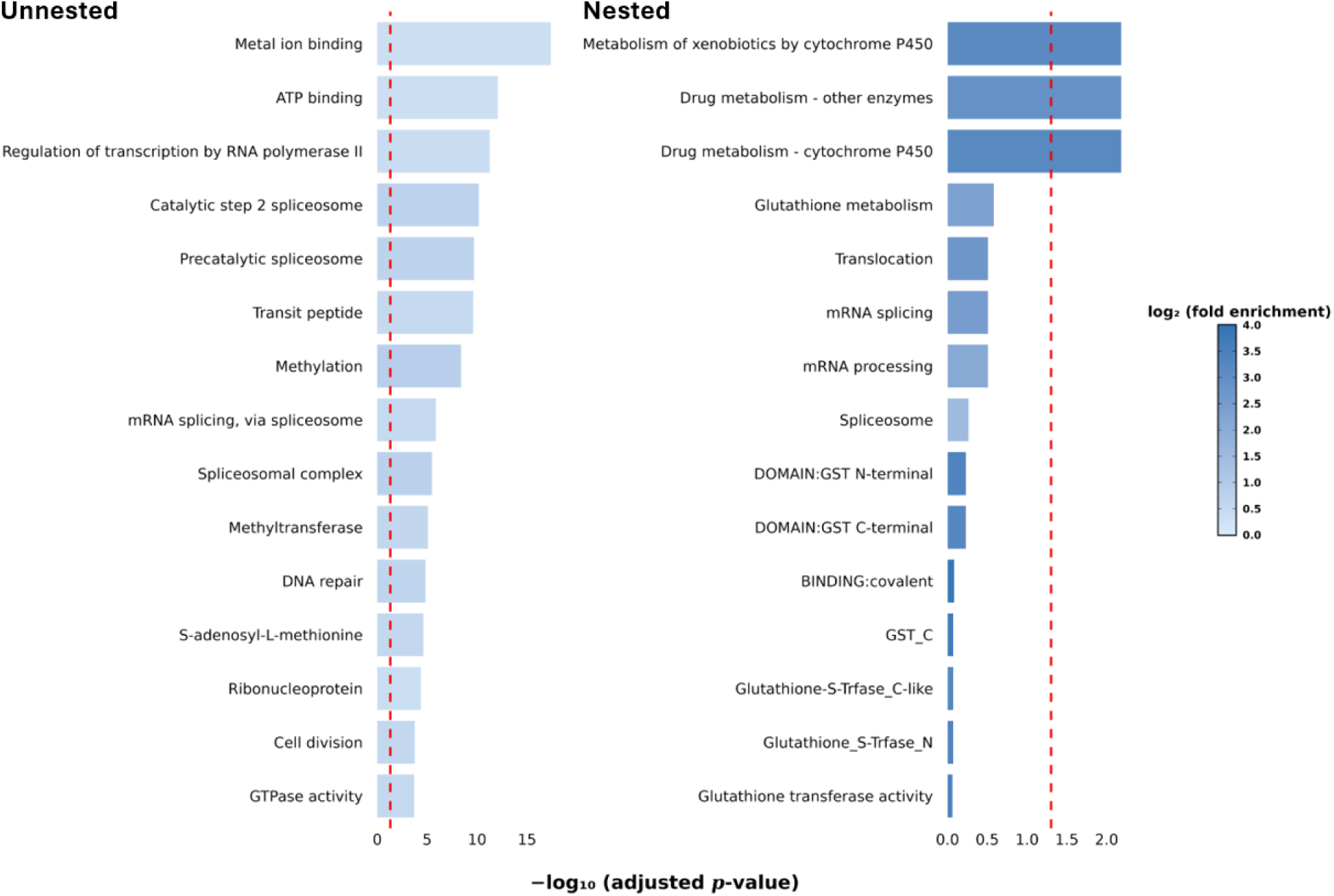
Visualization of the top 15 enriched terms for “diverged” *Drosophila* genes. Bar plots show terms ranked by Benjamini-adjusted *p*-values for unnested (left) and nested (right) genes, with the red dashed line indicating the threshold for statistical significance. The complete sets of enriched terms are provided in Tables S1 and S3.

To investigate functional themes associated with strong expression divergence, we examined the top-ranked unnested and nested ortholog pairs in each taxon, defined by the largest absolute differences in estimated expression optima. Among *Drosophila* unnested genes, the most divergent exemplar was Oligosaccharyltransferase Complex Subunit 4 (*Ost4*, FBgn0053774), a small ER transmembrane protein essential for N-linked glycosylation and protein maturation (Ruiz-Canada et al., 2009; Dumax-Vorzet et al., 2013; Katoh and Tiemeyer, 2013). The top mammalian unnested gene was Alpha-2-Macroglobulin (*A2M*, ENSG00000175899), a broad-spectrum protease inhibitor involved in cytokine transport, immune regulation, and proteostasis, with established links to inflammatory and neurodegenerative processes (Du et al., 1997; French et al., 2008; Wong and Dessen, 2014). Among *Drosophila* nested genes, the most divergent gene was Ubiquinol-Cytochrome c Reductase 11 kDa Subunit-Like (*UQCR-11L*, FBgn0050354), a mitochondrial complex III subunit previously identified as highly divergent in an analysis of a smaller set of relocated genes (Piya et al., 2023). In mammals, the top nested exemplar was Ribosomal Protein S27 (*RPS27*, ENSG00000177954), a zinc-binding component of the 40S ribosomal subunit involved in translation, rRNA processing, and stress response signaling that is frequently dysregulated in human disease (Zhang et al., 2003; Gazda et al., 2006; Xiong et al., 2014). Together, these cases highlight the distinct molecular functions associated with pronounced expression divergence and illustrate how genomic architecture can shape gene-specific evolutionary trajectories.

## Discussion

Our results show that broad taxonomic context and local genomic architecture leave distinct, interacting signatures on the evolution of gene expression across animals. Expression divergence was nearly four times more frequent in *Drosophila* than in mammals, consistent with expectations from population-genetic theory (Beckenbach et al., 1993; Lynch and Conery, 2003; Jensen and Bachtrog, 2011) and extensive evidence for rapid regulatory evolution in *Drosophila* (Britten, 1986; Moriyama, 1987; Carroll, 2005; Coolon et al., 2015; Nourmohammad et al., 2017). At the same time, genomic architecture added another layer of heterogeneity. In *Drosophila*, nested genes were less likely than unnested genes to diverge, yet those that did diverge displayed larger shifts in expression optima, with internal genes showing the strongest effects. In contrast, mammalian nested and unnested genes showed similar frequencies and magnitudes of expression divergence. Across both taxa, young nested genes diverged more frequently than old nested genes, suggesting a transient period of regulatory instability after nesting. However, the magnitudes of expression divergence did not differ by age, indicating that nesting does not typically initiate large expression shifts but instead shapes expression trajectories once divergence begins. Tissue-level analyses further revealed that divergence is distributed unevenly across organ systems in a taxon- and architecture-specific manner, and functional enrichment patterns mirrored these differences. Together, these findings illustrate that genome-wide patterns of expression divergence emerge from the interplay between lineage-specific biological constraints and the fine-scale physical organization of genomes.

The contrasting divergence patterns between *Drosophila* and mammals likely reflect long-standing differences in effective population size, regulatory complexity, and the strength of purifying selection (Beckenbach et al., 1993; Lynch and Conery, 2003; Jensen and Bachtrog, 2011). Regulatory evolution in *Drosophila* has been shown to proceed rapidly due to efficient selection and turnover of *cis*-regulatory elements (McGregor et al., 2007; Wittkopp and Kalay, 2012; Signor and Nuzhdin, 2018), whereas mammalian expression profiles are generally more constrained (Khaitovich et al., 2006; Brawand et al., 2011; Perry et al., 2012). Within each taxon, genomic architecture further structured divergence. The larger expression shifts observed among *Drosophila* nested genes—particularly internal genes—are consistent with hypotheses that transcriptional inteference, competition for regulatory elements, or altered chromatin states may affect expression dynamics of genes embedded within introns (Da Lage et al., 2003; Crampton et al., 2006; Osato et al., 2007; Assis et al., 2008). Elevated divergence rates among young nested genes in both taxa likewise parallel findings that relocated or recontextualized genes often experience rapid expression changes before stabilizing (Bhutkar et al., 2007; Kaessmann et al., 2009; Meisel, 2009; Egecioglu and Brickner, 2011; Assis and Bachtrog, 2013; Assis, 2016). Because divergence magnitudes did not differ by age, these results imply that architectural context influences how expression evolves after divergence begins rather than determining the extent of that divergence.

A key distinction between taxa emerges when considering internal genes. In *Drosophila*, the reduced overall prevalence of divergence among nested genes suggests that many internal genes arrive with expression profiles that are already sufficiently different from those of their external genes, reducing the likelihood that further divergence is necessary. However, when an internal gene does not arrive with an adequately differentiated regulatory profile, it can undergo rapid and pronounced divergence, resulting in the large shifts we observed. This pattern supports a model in which internal genes in *Drosophila* follow one of two trajectories: either they are inserted into introns with distinct expression profiles already in place, or they rapidly diverge to achieve them. In mammals, internal genes exhibited neither elevated divergence frequencies nor increased divergence magnitudes relative to external nested genes. These results suggest that mammalian internal genes must arrive with their regulatory distinctiveness already established, as they do not exhibit the rapid post-formation divergence trajectory observed in *Drosophila*. This contrast highlights fundamental differences in the evolutionary opportunities and constraints associated with genomic architectural changes across taxa.

Tissue-level analyses showed that expression divergence is distributed unevenly across organ systems in ways that differ between taxa and genomic contexts. These differences may reflect variation in tissue-specific regulatory constraints and functional demands. Among unnested genes, all three *Drosophila* reproductive tissues (testis, accessory gland, and ovary) showed consistently low divergence, whereas in mammals, testis was among the tissues showing the highest divergence. Relative to unnested genes, *Drosophila* nested genes exhibited approximately twofold higher divergence in testis and nearly 1.5-fold higher divergence in female and male head and accessory gland, suggesting that nested gene divergence within this taxon may be associated with reproductive and neural functions. Similarly, mammalian nested genes also displayed their highest divergence in brain and testis, in contrast to the elevated kidney divergence observed for unnested genes. Functional enrichment patterns further supported this view. Unnested genes were enriched for transcriptional regulation and RNA processing in *Drosophila* and for mitochondrial and membrane targeting in mammals, suggesting that different core cellular pathways contribute disproportionately to expression divergence in each taxon. *Drosophila* nested genes were enriched for detoxification and xenobiotic pathways, consistent with the rapid and lineage-specific evolution of environmental response genes (Thomas, 2007; Good et al., 2014; Dermauw et al., 2020; Darragh et al., 2021; Lu et al., 2021). In contrast, mammalian nested genes showed no clear functional enrichment, though top-ranked terms were associated with membrane signaling functions. These tissue-specific and functional differences underscore that taxonomic and architectural contexts together shape the regulatory pressures acting on individual genes.

Collectively, our findings motivate several directions for further investigation. First, expanding analyses to additional taxa, particularly those that differ in genome size, life history, and effective population size, would help clarify whether the architectural patterns observed here are lineage-specific or broadly conserved. Second, integrating regulatory sequence evolution, chromatin accessibility, and three-dimensional genome organization could elucidate mechanisms underlying the amplified expression divergence of internal nested genes, especially in light of emerging links between intragenic architecture and chromatin topologies (Bonev and Cavalli, 2016; Rowley and Corces, 2018). Third, finer-grained tissue resolution, especially in mammals, would improve power to detect subtle architectural effects and reveal whether particular cell types are especially prone to divergence in nested contexts. Finally, connecting expression divergence with phenotypic, ecological, or environmental data may illuminate cases in which architectural constraints or opportunities contribute to adaptive regulatory evolution. Together, such efforts will deepen our understanding of how taxonomic context and genomic architecture interact to shape the evolution of gene expression across animals.

## Methods

### Data collection and processing

Tables of quantile-normalized, log-transformed RNA-seq abundances measured in fragments per kilobase of transcript per million mapped fragments (FPKM) from carcass, female head, male head, ovary, testis, and accessory gland tissues in *Drosophila melanogaster* and *Drosophila pseudoobscura* were downloaded from the Dryad repository associated with Assis (2019) at https://datadryad.org/dataset/doi:10.5061/dryad.742564m. Analogous expression tables from brain, heart, lung, colon, liver, kidney, spleen, and testis tissues in human and mouse were downloaded from Expression Atlas (Kapushesky et al., 2010) at https://www.ebi.ac.uk/gxa/home/. Nested gene annotations were obtained from Assis (2016) for *Drosophila* and Assis (2021) for mammals. Tables of 1:1 orthologs were sourced from FlyBase (Öztürk Çolak et al., 2024) for *Drosophila* and Ensembl BioMart (Durinck et al., 2005, 2009) for mammals.

In total, there were 8,289 1:1 orthologs between *D. melanogaster* and *D. pseudoobscura* and 9,369 1:1 orthologs between human and mouse. Of these, 1,129 were nested in at least one *Drosophila* species, and 513 were nested in at least one mammal. In *Drosophila*, there were 659 internal and 470 external nested genes, reflecting the common occurrence of Russian-doll-like multi-level nesting (Assis et al., 2008; Yu et al., 2005). In mammals, there were 247 internal and 266 external nested genes, consistent with the lower overall frequency of nesting and the relative rarity of Russian-doll-like nested architectures in this taxon (Assis, 2021). Classifications of young and old nested genes were obtained from Assis (2016) for *Drosophila* and Assis (2021) for mammals. In *Drosophila*, 155 nested gene pairs were classified as young and 358 as old, whereas in mammals 384 nested gene pairs were young and 745 were old.

### Prediction of expression divergence and optima

The PiXi neural network (Piya et al., 2023) was used to predict expression divergence and species-specific expression optima for all orthologs in our study. For training, we generated 40,000 simulated observations, evenly divided between the two classes. Simulations followed the procedure of Piya et al. (2023), with OU parameters sampled from log_10_(*θ*) ∈ [0,5], log_10_(*α*) ∈ [0,3], and log_10_(*σ*^2^) ∈ [-2,3]. As in the original implementation Piya et al. (2023), we trained a two-layer neural network with a batch size of 5,000 for 500 epochs, with 25 values of *λ* uniformly sampled from log_10_(*λ*) ∈ [-12,-3] and *γ* ∈ {0, 0.1, , 1.0}. All 1:1 orthologs and their matched expression data (see *Acquisition of datasets used for analyses*) were provided as input to the trained PiXi model, which classified each ortholog pair as either “conserved” or “diverged” and predicted its expression optima (*θ*) in each species (Piya et al., 2023).

### Statistical analyses

All statistical analyses were performed in R (R Core Team, 2022) using the Posit Cloud IDE (RStudio Team, 2024). Two-tailed binomial tests were conducted with the binom.test() function (Abdi, 2007) to compare counts of *Drosophila* and mammalian “diverged” total, unnested, and nested genes, as well as counts of internal vs. external nested genes and young vs. old nested genes. For each test, we set the number of successes *X* to the number of genes in the focal category, the number of trials *n* to the total number of genes, and the null probability of success *p* to 0.5.

Two-tailed permutation tests, implemented with the perm_test() function in the stats package (R Core Team, 2022; RStudio Team, 2024), were used to evaluate differences between distributions shown in Figures 1 and 2. For each comparison, we calculated the absolute differences in *θ* across orthologous pairs and used the median of these values as the test statistic. Group labels were permuted 10,000 times to generate the null distribution, and empirical *p*-values were defined as the proportion of permuted statistics equalt to or more extreme than the observed statistic. For tissue-level analyses (Figure 2), *p*-values were Bonferroni-corrected for the number of tissues compared (six in *Drosophila* and eight in mammals) using the p.adjust() function in the stats package (R Core Team, 2022; RStudio Team, 2024).

### Functional enrichment analyses

We used the DAVID Functional Annotation Tool (Huang et al., 2009; Sherman et al., 2022) to evaluate biological functions associated with orthologs classified as “diverged” in each taxon. We performed two analyses for each taxon: one using “diverged” unnested genes as the target set and another using “diverged” nested genes as the target set. For all runs, we used the genome-wide background provided by DAVID. Enrichment was assessed across all annotation clusters available in DAVID. Statistical significance was determined using Fishers exact tests with BenjaminiHochberg FDR correction. This framework allowed us to identify whether diverged orthologs in unnested versus nested contexts were associated with particular biological processes or pathways.

## Data availability

All R code and processed datasets are available at https://github.com/PerryFAU/Nested_Genes.

## Acknowledgments

This work was supported by National Institutes of Health grant R35GM142438 and National Science Foundation grant DBI-2130666.

