## Supplement for "Taxonomic context and genomic architecture jointly shape expression divergence across animals"

Table S1: Functional enrichment results for “diverged” unnested genes in *Drosophila*.

| Source | Term | Adjusted $p$-value | Fold Enrichment |
|---|---|---|---|
| GOTERM_MF_DIRECT | Metal ion binding | 4.13E-18 | 1.30 |
| GOTERM_MF_DIRECT | ATP binding | 8.37E-13 | 1.30 |
| GOTERM_BP_DIRECT | Regulation of transcription by RNA polymerase II | 5.47E-12 | 1.33 |
| GOTERM_CC_DIRECT | Catalytic step 2 spliceosome | 6.58E-11 | 1.69 |
| GOTERM_CC_DIRECT | Precatalytic spliceosome | 2.16E-10 | 1.64 |
| UP_KW_DOMAIN | Transit peptide | 2.45E-10 | 1.46 |
| GOTERM_BP_DIRECT | Methylation | 4.14E-09 | 1.83 |
| GOTERM_BP_DIRECT | mRNA splicing, via spliceosome | 1.34E-06 | 1.43 |
| GOTERM_CC_DIRECT | Spliceosomal complex | 3.34E-06 | 1.74 |
| UP_KW_MOLECULAR_FUNCTION | Methyltransferase | 8.38E-06 | 1.52 |
| GOTERM_BP_DIRECT | DNA repair | 1.49E-05 | 1.59 |
| UP_KW_LIGAND | S-adenosyl-L-methionine | 2.39E-05 | 1.49 |
| UP_KW_MOLECULAR_FUNCTION | Ribonucleoprotein | 4.29E-05 | 1.31 |
| GOTERM_BP_DIRECT | Cell division | 1.71E-04 | 1.48 |
| GOTERM_MF_DIRECT | GTPase activity | 2.13E-04 | 1.45 |
| UP_KW_MOLECULAR_FUNCTION | Helicase | 2.27E-04 | 1.42 |
| UP_KW_MOLECULAR_FUNCTION | Kinase | 2.27E-04 | 1.22 |
| UP_KW_MOLECULAR_FUNCTION | Helicase | 2.27E-04 | 1.42 |
| GOTERM_MF_DIRECT | RNA polymerase II cis-regulatory region sequence-specific DNA binding | 5.63E-04 | 1.26 |
| UP_KW_CELLULAR_COMPONENT | Mitochondrion | 7.62E-04 | 1.16 |
| GOTERM_MF_DIRECT | Protein serine/threonine kinase activity | 1.00E-03 | 1.40 |
| GOTERM_MF_DIRECT | Protein serine/threonine kinase activity | 1.00E-03 | 1.40 |
| UP_SEQ_FEATURE | DOMAIN:C2H2-type | 1.38E-03 | 1.28 |
| GOTERM_BP_DIRECT | Mitochondrial respiratory chain complex I assembly | 1.59E-03 | 1.74 |
| SMART | ZnF_C2H2 | 1.82E-03 | 1.25 |
| SMART | SEC14 | 1.82E-03 | 1.80 |
| GOTERM_BP_DIRECT | Protein ubiquitination | 3.23E-03 | 1.36 |
| GOTERM_BP_DIRECT | Circadian rhythm | 3.23E-03 | 1.59 |
| GOTERM_BP_DIRECT | Transcription by RNA polymerase II | 3.28E-03 | 1.72 |
| UP_SEQ_FEATURE | TRANSIT:Mitochondrion | 3.43E-03 | 1.35 |
| UP_KW_DOMAIN | Kelch repeat | 3.59E-03 | 1.99 |
| GOTERM_MF_DIRECT | DNA-binding transcription factor activity, RNA polymerase II-specific | 3.88E-03 | 1.22 |
| GOTERM_MF_DIRECT | mRNA binding | 3.88E-03 | 1.31 |
| UP_KW_DOMAIN | WD repeat | 4.03E-03 | 1.33 |
| GOTERM_BP_DIRECT | DNA-templated DNA replication | 4.23E-03 | 1.95 |
| GOTERM_BP_DIRECT | Endosome transport via multivesicular body sorting pathway | 4.49E-03 | 2.12 |
| GOTERM_MF_DIRECT | Rho-dependent protein serine/threonine kinase activity | 4.87E-03 | 1.57 |
| GOTERM_MF_DIRECT | AMP-activated protein kinase activity | 4.87E-03 | 1.57 |
| GOTERM_MF_DIRECT | Eukaryotic translation initiation factor 2alpha kinase activity | 4.87E-03 | 1.57 |
| GOTERM_MF_DIRECT | Histone H3S28 kinase activity | 4.87E-03 | 1.56 |
| GOTERM_MF_DIRECT | Histone H4S1 kinase activity | 4.87E-03 | 1.56 |
| GOTERM_MF_DIRECT | Histone H2AS1 kinase activity | 4.87E-03 | 1.56 |
| GOTERM_MF_DIRECT | Histone H2BS14 kinase activity | 4.87E-03 | 1.56 |
| GOTERM_MF_DIRECT | Histone H3S57 kinase activity | 4.87E-03 | 1.56 |
| GOTERM_MF_DIRECT | Histone H3T45 kinase activity | 4.87E-03 | 1.56 |
| GOTERM_MF_DIRECT | Histone H2AXS139 kinase activity | 4.87E-03 | 1.56 |
| GOTERM_MF_DIRECT | Histone H3T3 kinase activity | 4.87E-03 | 1.56 |
| GOTERM_MF_DIRECT | Histone H2AS121 kinase activity | 4.87E-03 | 1.56 |
| GOTERM_MF_DIRECT | 3-phosphoinositide-dependent protein kinase activity | 4.87E-03 | 1.56 |
| GOTERM_MF_DIRECT | DNA-dependent protein kinase activity | 4.87E-03 | 1.56 |
| GOTERM_MF_DIRECT | Histone H3S10 kinase activity | 4.87E-03 | 1.56 |
| GOTERM_MF_DIRECT | Histone H2BS36 kinase activity | 4.87E-03 | 1.56 |
| GOTERM_MF_DIRECT | Ribosomal protein S6 kinase activity | 4.87E-03 | 1.56 |
| GOTERM_MF_DIRECT | Histone H3T11 kinase activity | 4.87E-03 | 1.56 |
| GOTERM_MF_DIRECT | Histone H3T6 kinase activity | 4.87E-03 | 1.56 |
| GOTERM_MF_DIRECT | Histone H2AT120 kinase activity | 4.87E-03 | 1.56 |
| GOTERM_BP_DIRECT | Mitochondrial translation | 4.98E-03 | 1.43 |
| INTERPRO | SAM-dependent_MTases_sf | 5.73E-03 | 1.48 |

| Source | Term | Adjusted *p*-value | Fold Enrichment |
|---|---|---|---|
| INTERPRO | Znf_C2H2_sf | 5.73E-03 | 1.26 |
| GOTERM_MF_DIRECT | GTP binding | 6.02E-03 | 1.31 |
| GOTERM_MF_DIRECT | Phosphatidylinositol bisphosphate binding | 7.55E-03 | 1.92 |
| GOTERM_MF_DIRECT | RNA helicase activity | 7.55E-03 | 1.61 |
| UP_KW_MOLECULAR_FUNCTION | Ribosomal protein | 8.31E-03 | 1.27 |
| UP_KW_DOMAIN | Leucine-rich repeat | 8.36E-03 | 1.31 |
| GOTERM_MF_DIRECT | Protein kinase activity | 1.01E-02 | 1.36 |
| UP_KW_DOMAIN | Transmembrane | 1.14E-02 | 1.04 |
| UP_KW_MOLECULAR_FUNCTION | Thiol protease | 1.21E-02 | 1.48 |
| UP_SEQ_FEATURE | DOMAIN:CRAL-TRIO | 1.27E-02 | 1.73 |
| UP_KW_CELLULAR_COMPONENT | Spliceosome | 1.32E-02 | 1.42 |
| GOTERM_MF_DIRECT | Ubiquitin-like ligase-substrate adaptor activity | 1.63E-02 | 1.81 |
| GOTERM_MF_DIRECT | SNARE binding | 1.71E-02 | 1.68 |
| GOTERM_MF_DIRECT | Protein serine kinase activity | 1.73E-02 | 1.40 |
| GOTERM_MF_DIRECT | Protein serine kinase activity | 1.73E-02 | 1.40 |
| GOTERM_BP_DIRECT | Transcription initiation at RNA polymerase II promoter | 2.69E-02 | 1.61 |
| GOTERM_MF_DIRECT | Four-way junction helicase activity | 2.81E-02 | 1.72 |
| GOTERM_CC_DIRECT | Vesicle | 2.84E-02 | 1.50 |
| UP_KW_MOLECULAR_FUNCTION | Serine/threonine-protein kinase | 3.01E-02 | 1.22 |
| UP_KW_MOLECULAR_FUNCTION | Serine/threonine-protein kinase | 3.01E-02 | 1.22 |
| GOTERM_CC_DIRECT | Cul3-RING ubiquitin ligase complex | 3.19E-02 | 2.00 |
| UP_KW_DOMAIN | ANK repeat | 3.46E-02 | 1.38 |
| UP_KW_BIOLOGICAL_PROCESS | Cell cycle | 3.50E-02 | 1.17 |
| GOTERM_MF_DIRECT | Transcription cis-regulatory region binding | 3.74E-02 | 1.37 |
| GOTERM_MF_DIRECT | DNA helicase activity | 3.75E-02 | 1.88 |
| GOTERM_MF_DIRECT | DNA clamp loader activity | 3.75E-02 | 1.47 |
| GOTERM_MF_DIRECT | Cysteine-type deubiquitinase activity | 3.91E-02 | 1.65 |
| GOTERM_MF_DIRECT | Single-stranded 3'-5' DNA helicase activity | 4.05E-02 | 1.70 |
| GOTERM_MF_DIRECT | Double-stranded DNA helicase activity | 4.05E-02 | 1.70 |
| INTERPRO | CRAL-TRIO_dom_sf | 4.28E-02 | 1.67 |
| GOTERM_MF_DIRECT | Pyridoxal phosphate binding | 4.62E-02 | 1.66 |

Table S2: Functional enrichment results for “diverged” unnested genes in mammals.

| Source | Term | Adjusted *p*-value | Fold Enrichment |
|---|---|---|---|
| GOTERM_CC_DIRECT | Mitochondrion | 3.54E-5 | 2.05 |
| UP_KW_CELLULAR_COMPONENT | Mitochondrion | 3.19E-4 | 1.96 |
| UP_KW_DOMAIN | Transit peptide | 3.31E-3 | 2.49 |
| GOTERM_CC_DIRECT | Membrane | 3.64E-3 | 1.36 |
| GOTERM_MF_DIRECT | S100 protein binding | 1.24E-2 | 19.76 |
| UP_KW_BIOLOGICAL_PROCESS | Ubl conjugation pathway | 2.47E-2 | 1.91 |
| GOTERM_MF_DIRECT | ATP binding | 4.97E-2 | 1.68 |

Table S3: Functional enrichment results for “diverged” nested genes in *Drosophila*.

| Source | Term | Adjusted *p*-value | Fold Enrichment |
|---|---|---|---|
| KEGG_PATHWAY | Drug metabolism - other enzymes | 6.57E-3 | 7.55 |
| KEGG_PATHWAY | Drug metabolism - cytochrome P450 | 6.57E-3 | 9.14 |
| KEGG_PATHWAY | Metabolism of xenobiotics by cytochrome P450 | 6.57E-3 | 8.88 |
